# Light on its feet: Acclimation to high and low diurnal light is flexible in *Chlamydomonas reinhardtii*

**DOI:** 10.1101/2025.08.28.672467

**Authors:** Sunnyjoy Dupuis, Jordan L. Chastain, Genevieve Han, Victor Zhong, Sean D. Gallaher, Carrie D. Nicora, Samuel O. Purvine, Mary S. Lipton, Krishna K. Niyogi, Masakazu Iwai, Sabeeha S. Merchant

## Abstract

Chlamydomonas acclimates to repeated low (LL) or high light (HL) days by changing the abundance of photosynthetic complexes and the ultrastructure of its thylakoid membranes. These phenotypes persist through the night phases, suggesting a readiness for the daylight environment that is routinely experienced despite the intervening dark periods (Dupuis & Ojeda *et al*. 2025). Here, we investigate how prior acclimation impacts algal fitness upon a change in daylight intensity and how quickly Chlamydomonas can reprogram its photoprotective strategy in a diurnal context. We performed a systems analysis of synchronized populations acclimated to diurnal LL when subjected to HL days and of populations acclimated to diurnal HL when subjected to LL days. In the latter case, diurnal photoacclimation decreased fitness during the first day at a new light intensity: HL-acclimated cells barely increased in size over the first LL period, and they failed to complete a cell cycle. However, although LL-acclimated cells showed severe photodamage after 6 hours of HL, they recovered chloroplast form and function later that afternoon and successfully divided at nightfall. These cells rapidly altered their thylakoid membrane ultrastructure, increased their photoprotective quenching capacity, and decreased their inventory of photosystem and antenna proteins by the end of the first HL day. Transcriptomic and proteomic analyses revealed rapid induction of thousands of genes, including those encoding proteases, chaperones, and other proteins involved in the chloroplast unfolded protein response. These results show that the alga is highly flexible and competent to rapidly acclimate to changes in diurnal light intensity.

## Introduction

Algae are beholden to the Sun and the dynamic light environments found on our planet. As the Sun rises, sets, and is blocked by clouds or other obstructions, the incident light intensity can vary by several orders of magnitude (1–3). Algae must absorb the incident light with light-harvesting pigment-protein complexes and direct the energy to the reaction centers of their photosystems to generate chemical energy for carbon assimilation and growth. Yet, absorbing more light energy than can be processed by the slower downstream reactions can cause photooxidative damage and decrease photosynthetic efficiency.

To succeed in dynamic light environments, photosynthetic microbes modulate the amount of light they absorb, dissipate excess light as heat through nonphotochemical quenching (NPQ), and repair photooxidative damage. While NPQ mechanisms can be activated and relaxed within minutes (4, 5), changes in chlorophyll (Chl) content and light-harvesting antenna size occur only after hours of exposure to high light (HL) or low light (LL), a process called photoacclimation (6–9). Upon prolonged HL, the popular reference alga *Chlamydomonas reinhardtii* (Chlamydomonas hereafter) decreases its cellular Chl content and light-harvesting complex (LHC) abundance to reduce the excitation energy pressure on the photosystems (10–15). In addition, the thylakoid membranes that house the LHCs and photosystems undergo changes in surface area and stacking, altering the interactions between the photosynthetic electron transfer chain complexes and other proteins, including those important for photosystem II (PSII) repair (6, 16, 17).

Prolonged HL exposure also leads to accumulation of the stress-related LHC-like proteins LHCSR3 and LHCSR1, which perform energy-dependent NPQ (qE) in Chlamydomonas (18–20). Upon acidification of the thylakoid lumen in excess light, these Chl- and carotenoid-binding proteins undergo rapid, pH-dependent conformational changes to turn into strong quenchers of excited Chl (19, 21–25). Expression of *LHCSR* and *PSBS* transcripts is suppressed in the absence of excess light or other activating signals (3, 26–29), as even their unprotonated protein products can quench excited Chl and decrease photosynthetic efficiency in LL (22). Yet, LHCSR proteins exhibit long half-lives in HL-acclimated Chlamydomonas cells transitioned to LL (20–30 h), persisting over the diurnal cycle and leading to constitutive NPQ capacity (30–32).

Several studies have investigated the kinetics of photoacclimation in Chlamydomonas transitioned between continuous LL and HL (11, 13, 33–36). These studies suggest that the alga can begin decreasing cellular Chl content, LHC abundance, and thylakoid membrane stacking within just 1–2 h of increased irradiance. However, the diurnal light cycles of our natural environment lead to rhythmic growth, metabolism, and gene expression, which could impact acclimation competency at a given time of day. In addition, the periodic darkness of night may interrupt acclimation processes and delay changes in cellular Chl content (37).

Chlamydomonas is an excellent reference organism for studying diurnal behavior. Like other photosynthetic organisms, its growth and metabolism are coordinated with time of day thanks to the periodic availability of light, which serves as both an energy source and a signal (38). As a result, Chlamydomonas populations synchronize when grown under repeated diurnal cycles in the laboratory: cells progress through the G1 phase during the daytime, undergo multiple fission S/M cycles at dusk to produce daughter cells of equal sizes, and then remain in a quiescent G0 phase of the cell cycle during the night. Synchronous populations provide high signal-to-noise in measurements of physiology and gene expression (31).

Recently, we documented the physiology and gene expression profile of synchronized Chlamydomonas populations acclimated to diurnal cycles of LL, moderate light (ML), and HL (50, 200, and 1000 µmol photons m^−2^ s^−1^, respectively) (30). We showed that photoacclimatory phenotypes persist in the night phase. Even after 10 h of darkness, populations acclimated to diurnal LL exhibited high Chl content, LHC protein abundance, and tightly stacked thylakoid membranes, while HL-acclimated populations maintained relatively loose stacks of thylakoid membranes, low photosystem and antenna protein abundance, and high NPQ capacity. We hypothesized that maintenance of this physiological state may be beneficial for cells that routinely encounter dim days or bright days, but that it likely decreases fitness upon a change in daylight intensity. To test this hypothesis, we transitioned LL-acclimated populations to diurnal HL and HL-acclimated populations to diurnal LL. We show that Chlamydomonas is flexible and can reacclimate in just one day despite severe photoinhibition. Using transcriptomics and proteomics, we document the gene expression program for re-acclimation with high temporal resolution over two diurnal cycles.

## Results

### LL-acclimated cells achieve impressive growth and recovery during their first day in HL despite severe photoinhibition, while acclimation to HL stunts growth upon a transition to diurnal LL

To determine the kinetics of diurnal photoacclimation in Chlamydomonas, we acclimated populations to diurnal LL or HL using the conditions established in the previous study (Materials and Methods) (30), and then surprised them with the opposite light (LL to HL, and HL to LL). We monitored the populations for two consecutive diurnal cycles, diluting the populations to the starting density after the first day (Figure S1). To test whether acclimation to diurnal LL decreased fitness in diurnal HL, we monitored photosynthetic efficiency and growth. The maximum efficiency of PSII (F_v_/F_m_) of the transitioned cells plummeted from 0.67±0.02 prior to the transition to 0.16±0.07 in the middle of the first HL day, indicating severe photoinhibition (Figure 1A) (30). Then, we observed an impressive recovery in the latter half of the day. By the night phase, the F_v_/F_m_ had rebounded to 0.61±0.04 and it then tracked closely with that of populations that had been acclimated to diurnal HL for several weeks.

**Figure 1.**
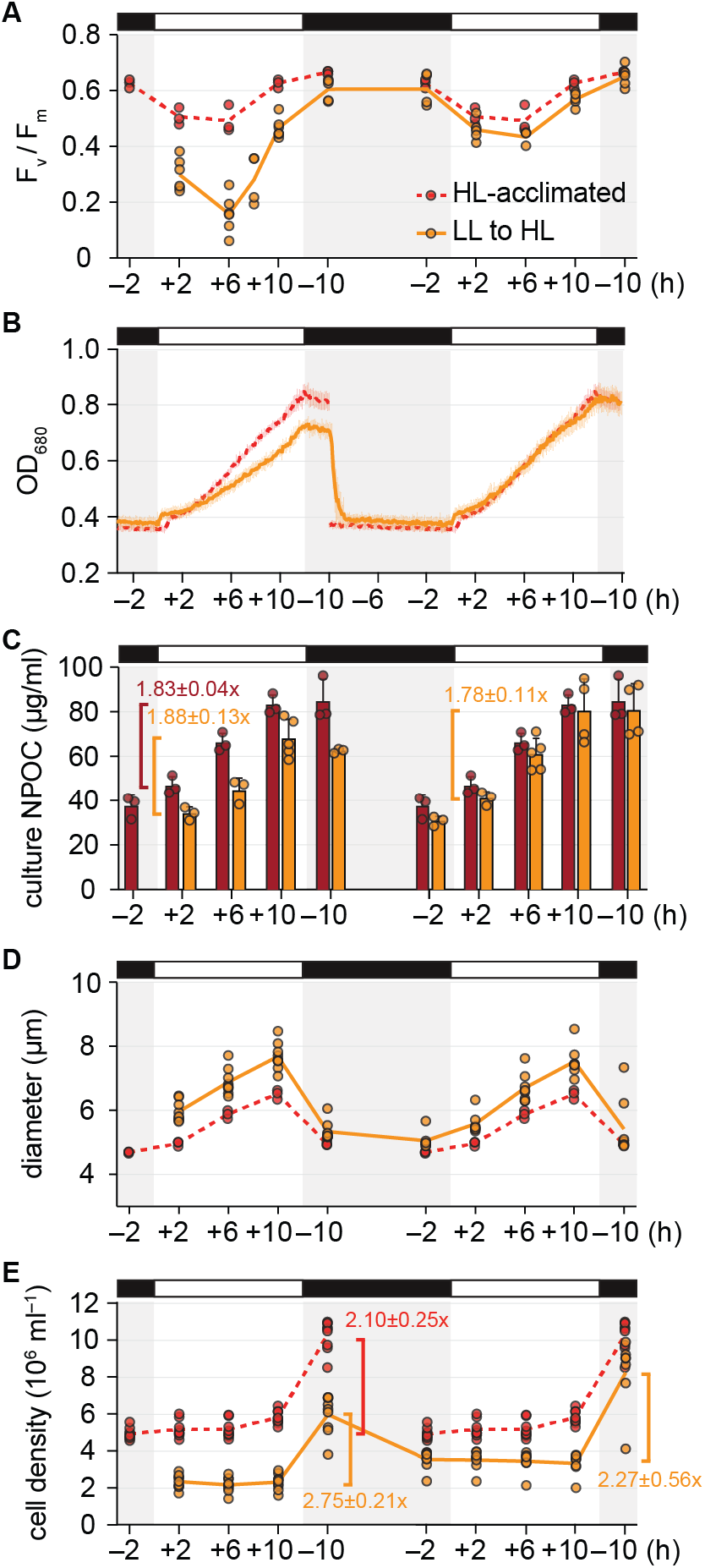
LL-acclimated cells achieve impressive growth and recovery during their first day in HL despite severe photoinhibition. The physiology of LL-acclimated cells transitioned to diurnal HL (orange) was monitored for two diurnal cycles. Results for HL-acclimated populations (red) are reproduced from Dupuis & Ojeda *et al*., 2025. (A) Maximum photochemical efficiency of PSII (F_v_/F_m_). (B) OD_680_ monitored continuously. Error bars represent the standard deviation from the mean. (C) NPOC concentration. Error bars represent the standard deviation from the mean. Fold-changes during the light phase are noted above the brackets as mean±standard deviation. (D) Mean cell diameter. (E) Cell density. Fold-changes over the day are noted near the brackets as mean±standard deviation.

We found that optical density (OD_680_) increased less for LL-acclimated populations during a surprise HL day than for populations acclimated to diurnal HL (Figure 1B). Yet, by just their second day in HL, the previously LL-acclimated cultures achieved the same increase in OD_680_ as did HL-acclimated cultures. Since Chl content may influence OD_680_, we measured nonpurgeable organic carbon (NPOC) content as another proxy for biomass. Although all cultures were adjusted to an OD_680_ near 0.4 at the beginning of the experiment, the starting NPOC content of transitioned cultures was lower than that of acclimated cultures (Figure 1C). However, the fold-change in the NPOC content of transitioned cultures over the first HL day was similar to that of HL-acclimated cultures. Thus, LL-acclimated cells could achieve similar increases in biomass over a surprise HL day despite the severe photoinhibition.

LL-acclimated cells were larger than the HL-acclimated cells across the two-day time course, particularly prior to the first division event (Figure 1D). The LL-to-HL transitioned cells successfully divided at the end of their first day in HL, and in fact, achieved a greater fold-change in cell density than did the HL-acclimated cells (Figure 1E). These results suggest that although acclimation to LL causes severe photoinhibition during a surprise HL day, it results in greater vegetative growth of Chlamydomonas than is achieved by cells that are long-term acclimated to diurnal HL.

We also determined how acclimation to diurnal HL influenced Chlamydomonas’ fitness upon a transition to diurnal LL. HL-acclimated populations exhibited high F_v_/F_m_ in LL and achieved similar increases in OD_680_ and NPOC content as did LL-acclimated populations (Figures S2A– S2C). However, the HL-to-LL transitioned cells were unable to sufficiently increase their size and hence failed to divide after their first day in LL (Figures S2D–S2E). Thus, acclimation to diurnal HL limits growth during LL days, decreasing fitness upon a change in the diurnal light environment. By the second day, the HL-to-LL cells became larger like LL-acclimated cells did, and some cells successfully divided at the light-to-dark transition.

### LL-acclimated cells alter their thylakoid membrane architecture and exhibit signs of chloroplast swelling upon a transition to HL

Components of the PSII repair pathway (e.g., FTSH protease, STL1 kinase, chloroplast ribosomes) are sterically excluded from appressed membrane regions where PSII is localized (17, 39, 40). Decreased stacking of appressed thylakoid membranes is thought to be important for PSII repair in prolonged HL (17, 41). Stacking is driven by electrostatic forces and van der Waals interactions between the lipids and proteins of adjacent membranes (42). Thus, changes in LHCII abundance, their phosphorylation by STT7 kinase, and the osmolarity of the chloroplast stroma and thylakoid lumen can alter thylakoid stacking (35, 43–45). Therefore, we tested whether the recovery in F_v_/F_m_ observed in the LL-to-HL transition might be accompanied by changes in thylakoid membrane stacking.

We obtained subdiffraction-resolution micrographs of Chl autofluorescence in live Chlamydomonas cells during the transition using confocal Airyscan microscopy. At the beginning of the HL day, the chloroplasts of LL-acclimated cells contained thick regions of Chl throughout the lobes and base, as has been reported previously for LL-acclimated Chlamydomonas (Figure 2A) (30, 35). Then, by the first +6 timepoint, the chloroplast displayed discrete, thin structures of Chl fluorescence, suggesting a substantial decrease in thylakoid membrane stacking. During the second day in HL, the fluorescent structures appeared less dispersed than they had in the first HL afternoon but still appeared thinner than at the start of the transition.

**Figure 2.**
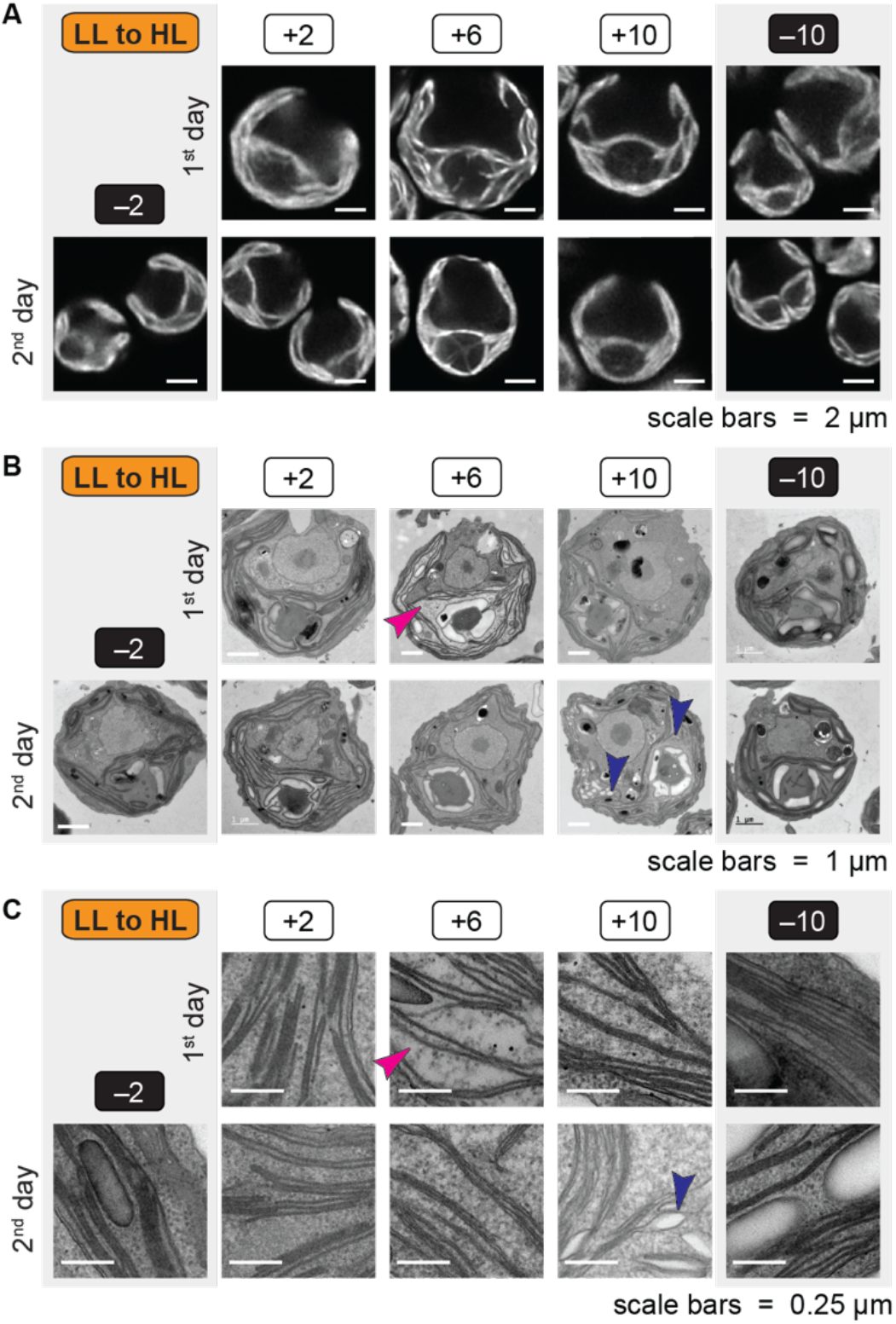
LL-acclimated cells rapidly alter their thylakoid membrane architecture and exhibit signs of chloroplast swelling upon a transition to HL. (A) Chl autofluorescence of representative live cells during the transition from diurnal LL to HL imaged by confocal Airyscan microscopy. Each Chlamydomonas cell has a single cup-shaped chloroplast with a pyrenoid at its base. (B) Representative electron micrographs of fixed cells during the transition from diurnal LL to HL. The pink arrowhead indicates the light stroma phenotype. Indigo arrowheads indicate examples of swollen thylakoids. (C) Representative electron micrographs of thylakoid membranes, with notable phenotypes indicated as in (B).

As a complementary approach, we fixed cells at each timepoint during the LL-to-HL transition and imaged them by transmission electron microscopy (TEM). The micrographs confirmed that thylakoid membrane stacking decreased within the first day of HL (Figures 2B–2C). Stacking was particularly low at the first +6 timepoint when we observed the lowest F_v_/F_m_ (Figure 1A): the thylakoid membranes were often present as single or double layers at this time. Across the population, the stroma of the chloroplast appeared less dense with ribosomes and other large densities at this timepoint (Figure 2B–2C, pink arrowheads). Such densities were strikingly dispersed from one another relative to those of the cytosol, which appeared more crowded than at other timepoints. This suggests that the entire chloroplast becomes swollen at this timepoint of the transition. Chloroplast swelling has been documented in Arabidopsis during HL stress as a result of damage to the chloroplast envelope and changes in osmolarity (46, 47). Changes in the osmolarity of the stroma could impact thylakoid membrane appression (48, 49). The prevalent “light stroma” phenotype became less evident in the population just 4 h later and was not observed during the second day of the transition.

Besides changes in stacking and the appearance of the stroma, we also observed swollen thylakoids in some cells of the LL-to-HL population, particularly at the +10 timepoints (Figure 2B– 2C, indigo arrowheads). Thylakoid swelling has been reported in Chlamydomonas exposed to photoinhibitory light intensities and has been attributed to increased accumulation of ammonium ions in the thylakoid lumen upon acidification and cyclic electron flow (50–52).

### Global view of gene expression reveals widespread mRNA induction at the onset of a surprising HL day

To understand the molecular events underlying the recovery in photosynthesis and chloroplast integrity, and to further explore photoacclimation processes in Chlamydomonas, we monitored the transcriptome and proteome of the LL- and HL-acclimated populations as they transitioned to a new intensity of diurnal light (Datasets S1 and S2). We began collecting RNA samples 30 min after lights on (+0.5), as we anticipated a rapid transcriptional response to a surprising quantity of light. Principal component analysis (PCA) of the transcriptome over the two-day LL-to-HL transition separated the data primarily by time of day, as observed previously for LL- and HL-acclimated Chlamydomonas (Figure 3A) (30). Samples collected at the same times from the first day (triangles) and second day (squares) in HL were mostly positioned near each other. However, the transcriptome at the first +0.5 and +1 timepoints was quite distinct from the same times on the second HL day, as well as from the subsequent timepoint (first +2) on PC1, which represented half of the variation in the data. No such distinction between these timepoints was apparent in PC1 of the HL-to-LL transition, which represented 85% of the variation in the transcriptome (Figure 3B).

**Figure 3.**
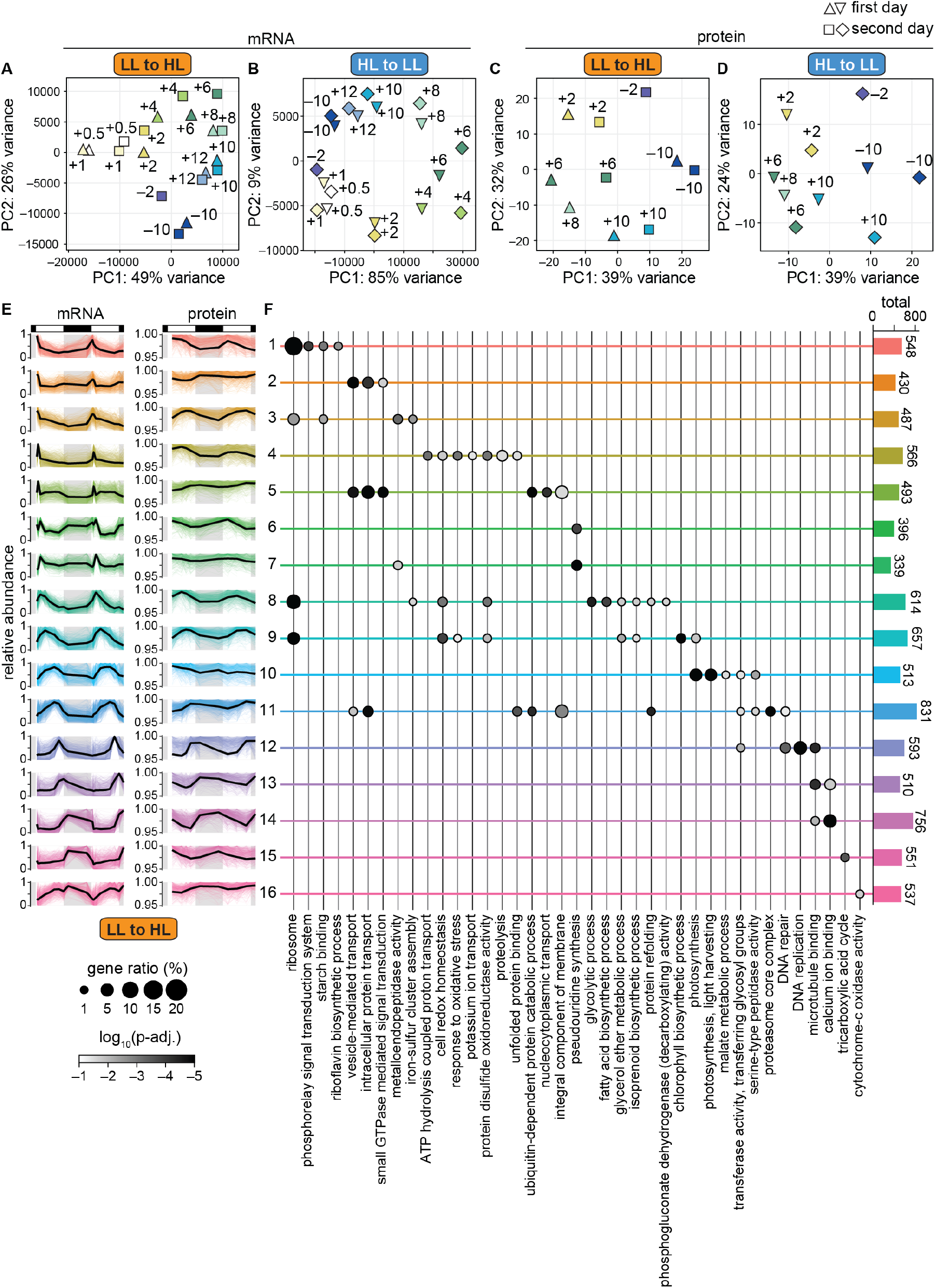
Global view of gene expression reveals maintenance of diurnal program, as well as widespread mRNA induction at the onset of a surprising HL day. (A) PCA of the transcriptome of cells transitioned from diurnal LL to HL at 19 timepoints. Samples collected on the first day are indicated as triangles, and those collected on the second day as squares. (B) PCA of the transcriptome of cells transitioned from diurnal HL to LL at 19 timepoints. Samples collected on the first day are indicated as triangles, and those collected on the second day as diamonds. (C) PCA of the proteome of cells transitioned from diurnal LL to HL at 10 timepoints, represented as in (A). (D) PCA of the proteome of cells transitioned from diurnal HL to LL as in (B). (E) Pattern of mRNA (left) and protein (right) accumulation for 16 distinct clusters of co-expressing genes during the LL-to-HL transition. Abundance was normalized relative to the maximum for each gene during the LL-to-HL transition. Colored lines show the relative abundance of individual mRNAs and proteins, while the black line shows the mean relative abundance. Cluster assignment is available in Dataset S2. (F) Enrichment of representative GO terms in the 16 clusters of genes. Bars at the right show the total number of genes in each cluster. The full list of significantly enriched GO terms is available in Dataset S3.

Time of day also explained the majority of the variation in the proteome for both transitions (Figures 3C–3D), as observed previously (30). Of the timepoints when the proteome was examined, the +6 timepoint appeared the most distinct between the first and second day in HL.

We investigated the changes in mRNA abundances that occur in LL-acclimated cells at the onset of the first HL day and how these changes shape the proteome. We detected 8,821 cognate mRNAs and proteins across the full time course and grouped these into 16 clusters based on how their relative abundances changed over time (Figure 3E, Dataset S2). mRNAs in many of the clusters exhibited a sharp peak in abundance within the first 2 h of HL (Clusters 4–7, ∼20% of the mRNAs included in the analysis). mRNAs in these clusters often exhibited a similar abundance peak during the second HL day, though its magnitude often differed from that of the first HL day. This induction was not evident in any groups of coexpressing mRNAs during the onset of the HL- to-LL transition (Figure S3, Dataset S2). However, many mRNAs did exhibit such an induction at lights on during the second day of the HL-to-LL transition.

Among the mRNAs that peak within the first 2 h of the LL-to-HL transition were genes enriched for Gene Ontology (GO) terms related to oxidative stress, ion transport, and proteolysis (Cluster 4 genes, whose proteins were most abundant midday) (Figures 3E–3F, Dataset S3). Genes in Cluster 5, whose proteins accumulated in the latter half of the day, were enriched for vesicular transport, protein transport, and ubiquitin related GO terms. Clusters 6 and 7 were enriched for terms related to metalloendopeptidases, transcription, and RNA modification. Thus, the mRNA induction at the start of the transition supports redox homeostasis, proteostasis, and housekeeping functions.

This global view of the gene expression landscape also revealed adjustments to the proteome in the second day of the LL-to-HL transition relative to the first. Proteins in Clusters 1, 4, 8, and 10 often appeared less abundant the second day, and were enriched for photosynthesis and light harvesting GO terms (Figures 3E–3F, Dataset S3). Proteins in Clusters 5 and 11 often appeared more abundant in the second day in HL, and were enriched for functions in proteostasis.

Taken together, we find that Chlamydomonas’ rhythmic gene expression program is maintained in severe light stress and that the surprise of excess light can induce widespread transcriptional changes that may be important for restoring homeostasis after photodamage without disrupting the diurnal program.

### NPQ mechanisms are activated within the first HL day

Unsurprisingly, genes encoding photoprotective functions were among the genes that exhibited mRNA induction at the onset of the transition from diurnal LL to HL. The expression of qE-related genes *LHCSRs* and *PSBSs* was previously reported to peak at lights on in diurnal ML and LL (200 and 60 µmol photons m^−2^ s^−1^, respectively), with the magnitude of expression being correlated to the intensity of light (31). We found that the induction of the *LHCSRs* was much higher during lights on at the first HL period than at the second (Figure 4A). Large increases in mRNA abundance at the first instance of HL may enable *de novo* accumulation of LHCSR3 and LHCSR1 proteins in the LL-acclimated cells, which steadily accumulated over the first HL day (Figure 4B). Conversely, HL-acclimated cells transitioned to LL did not induce *LHCSRs* during the first LL morning, but did during the second (Figure 4C), and LHCSR protein abundance decreased over the first diurnal cycle of LL (Figure 4D).

**Figure 4.**
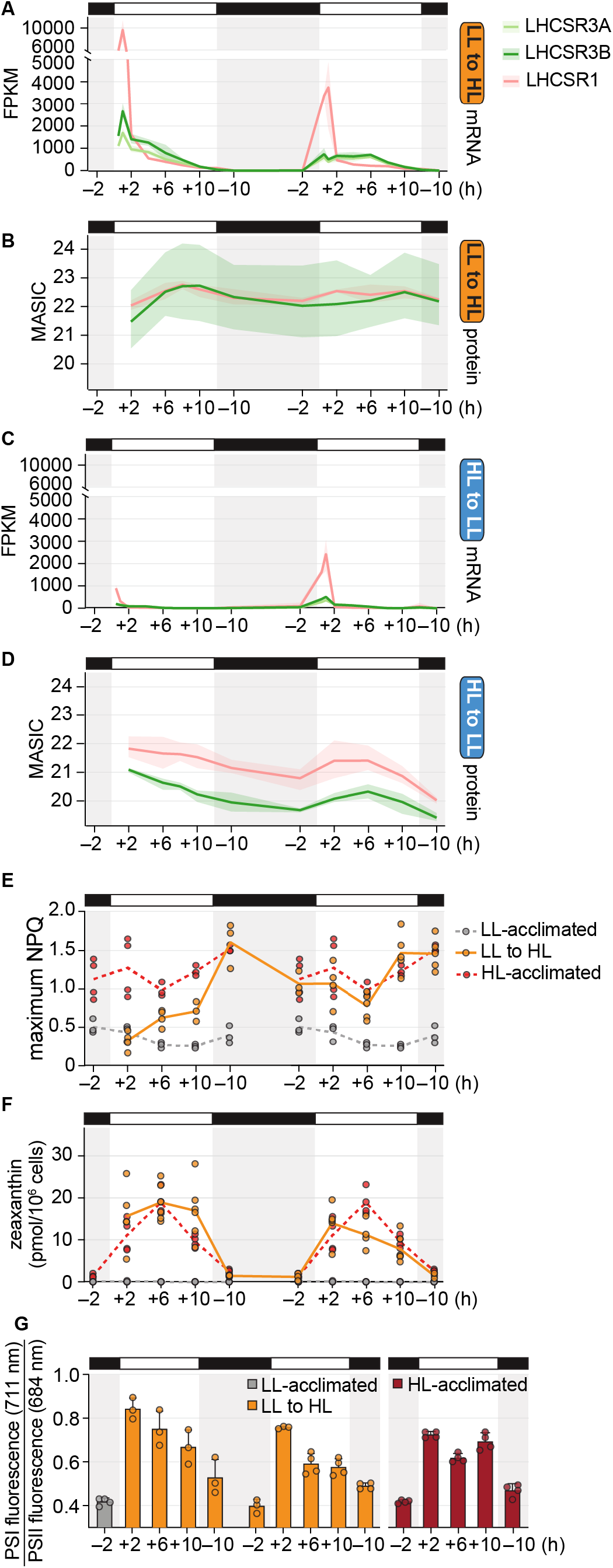
NPQ mechanisms are activated within the first HL day. Gene expression data are represented as the mean and the 95% confidence interval. Results for HL-acclimated populations (red) and LL-acclimated populations (grey) are reproduced from Dupuis & Ojeda *et al*., 2025. (A) Abundance of LHCSR transcripts during the diurnal LL-to-HL transition. (B) Abundance of LHCSR proteins during the diurnal LL-to-HL transition. (C) Abundance of LHCSR transcripts during the diurnal HL-to-LL transition. (D) Abundance of LHCSR proteins during the diurnal HL-to-LL transition. (E) Maximum NPQ measured over 50–1500 µmol photons m^−2^ s^−1^ at each timepoint. (F) Cellular zeaxanthin content. (G) Relative Chl fluorescence from PSI (at 711 nm) to PSII (at 684 nm) measured at 77 K.

To determine how these gene expression changes influenced photoprotective quenching, we first monitored NPQ capacity during the transitions. In both cases, NPQ capacity was correlated with LHCSR accumulation. LL-acclimated cells increased their NPQ capacity from 0.31±0.12 at the first +2 timepoint in HL to 0.70±0.13 at the first +10, and by the dark phase, had reached the capacity of cells that had been acclimated to HL for several weeks (1.60±0.30) (Figure 4E). Conversely, the NPQ capacity of HL-acclimated cells transitioned to LL decreased as LHCSR abundance decreased over the first day (Figure S4A).

Upon HL exposure, a high ΔpH across the thylakoid membrane activates CVDE1, which converts violaxanthin to the quenching pigment zeaxanthin. HPLC measurements showed that LL-acclimated cells were competent to accumulate zeaxanthin to the same degree as HL-acclimated cells do by the first +2 in HL (Figure 4F). Their de-epoxidation state followed a similar pattern as in the HL-acclimated cells (Figure S4C). HL-acclimated cells were also competent to adjust zeaxanthin levels to LL-acclimated levels by the first +2 of a surprising LL day (Figures S4B–S4C).

We also investigated state transitions in the populations by measuring Chl fluorescence spectra at 77 K. Within minutes of HL, STT7 kinase is known to phosphorylate LHCII trimers, increasing their association with PSI and thereby redistributing excitation energy between PSI and PSII (53, 54). We found that while LL-acclimated cells maintained the relative fluorescence of PSI:PSII near 0.4 at the beginning of the LL day (30), cells transitioned to HL doubled their relative PSI:PSII fluorescence to 0.89±0.05 within the first 2 h (Figure 4G). This increase was reversible, decreasing again in the night phase as it does for HL-acclimated cells. Such increases in PSI:PSII fluorescence may reflect not only association of LHCII with PSI, but also PSII damage.

In summary, these data suggest that LL-acclimated cells are competent to perform zeaxanthin-dependent quenching (qZ) and state transition-dependent quenching (qT) within just 2 h of a HL day. After several more hours, they accumulate LHCSR protein and increase their overall NPQ capacity, which may be aided by superinduction of *LHCSR* mRNAs at lights on of the transition. This gradual increase in overall NPQ capacity may contribute to the observed recovery in F_v_/F_m_ (Figure 1A).

### Transient depletion and delayed accumulation of photosystem complex transcripts may mediate the reduction in light harvesting during acclimation to diurnal HL

Next, we sought to determine how Chlamydomonas’ light-harvesting capacity and photosystem abundance are adjusted during the diurnal transitions. HPLC measurements showed that the Chl content of LL-acclimated cells transitioned to HL was higher than that of the HL-acclimated cells during the first HL day (Figure 5A). After the cells divided (Figure 1), their Chl content became more similar to that of HL-acclimated cells, and throughout the second HL day, it was lower than it had been the previous day. Conversely, HL-acclimated cells had lower Chl content than LL-acclimated cells had during their first day in LL (Figure 5B). As the HL-to-LL transitioned cells failed to divide after the first LL day and remained relatively large during the dark phase (Figure S2), they began the second LL day with a higher cellular Chl content than the LL-acclimated cells did. They continued to increase their Chl content over the second day of the transition.

**Figure 5.**
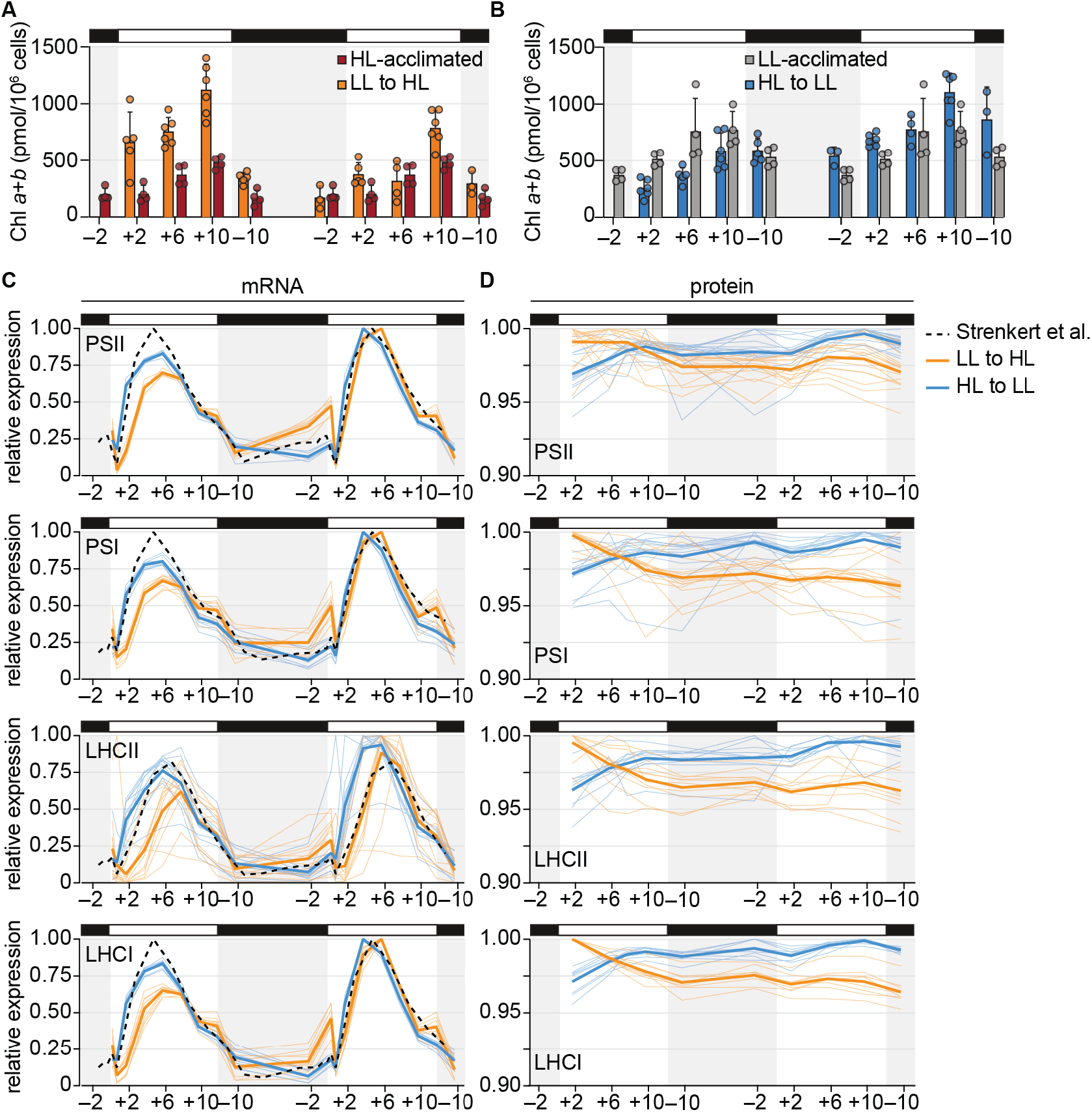
Transient depletion and delayed accumulation of photosystem complex transcripts may reduce absorptive capacity during acclimation to diurnal HL. Results for HL-acclimated populations (red) and LL-acclimated populations (grey) are reproduced from Dupuis & Ojeda *et al*., 2025. Mean mRNA and protein abundances were normalized relative to the maximum observed in each population (LL to HL normalized relative to the maximum observed in LL to HL; HL to LL normalized relative to the maximum observed in HL to LL). Fine lines show the relative abundance of individual mRNAs and proteins as the mean across the three experimental replicates, and the thick lines show the mean across all detected mRNAs or proteins of the photosystems and their antenna. (A) Cellular Chl content during the diurnal LL-to-HL transition. (B) Cellular Chl content during the diurnal HL-to-LL transition. (C) Changes in photosystem and antenna mRNA abundances during the diurnal transitions compared to that of ML-acclimated cells reported by Strenkert *et al*., 2019. (D) Changes in photosystem and antenna protein abundances during the diurnal transitions.

We found that transcripts for both photosystems and their respective LHCs sharply decreased in abundance during the first hours of the LL-to-HL transition (Figure 5C). A slight trough in PSII and LHCII mRNA abundance can also be observed in cells acclimated to ML (31) and in the HL-acclimated cells on the second LL day. In addition, mRNA accumulation was delayed during the first day of the LL-to-HL transition relative to diurnal ML conditions, whereas they accumulated on time or even earlier during the HL-to-LL transition.

The abundance of the cognate PSI, LHCII, and LHCI proteins showed a concerted decrease over the first day of the LL-to-HL transition (Figure 5D). The abundance of PSII subunits often did not begin to decrease until the latter half of the first day in HL, consistent with reports that translation of some PSII subunits (i.e., D1, encoded by *psbA*) transiently increases upon HL exposure to support PSII repair (11, 55, 56). Conversely, the abundance of the photosystem and antenna proteins increased over the HL-to-LL transition (Figure 5D). Collectively, these results suggest that changes in photosystem and antenna abundance are likely driven by changes not only in synthesis but also in degradation.

### Populations acclimated to diurnal LL promptly induce chaperones, proteases, and the autophagy pathway during a surprise HL morning

To distinguish degradation events during the transition from diurnal LL to HL, we investigated the expression of chloroplast-localized chaperones, proteases, and their regulators. LL-acclimated cells had the transcriptional signature of a heat shock or unfolded protein response during the first hour of HL (Figure 6A). Target loci of the heat shock transcription factor HSF1, including the *HSF1* locus itself, loci of the cytosolic chaperones *HSP90A* and *HSP70A*, and loci of the chloroplast chaperones *HSP22C, HSP22E, HSP22F*, and *HSP70B* exhibited high mRNA abundances at the +1 timepoint, particularly during the first day of the transition (Figure 6A) (57, 58). The same was true for several cochaperones and HSP100s, the *MARS1* kinase, which has been proposed to regulate the chloroplast unfolded protein response through retrograde signaling (59), and the VIPP paralogs, which are thought to play a role in the response to chloroplast membrane stress (52, 60). In addition, transcripts encoding the soluble chloroplast serine endopeptidases DEG1A and DEG1C and subunits of the thylakoid-membrane-tethered processive metalloprotease FTSH1 and FTSH2, which degrade the D1 polypeptide during PSII repair (61, 62), were also increased within the first hour of the transition. Transcriptional induction of several of these genes has been reported previously upon a dark-to-light transition or under continuous HL (59, 60, 62–64). However, the level of induction measured in the LL-to-HL population was not observed in the HL-to-LL transition (Figure S5) nor in the ML-acclimated populations studied previously (31).

**Figure 6.**
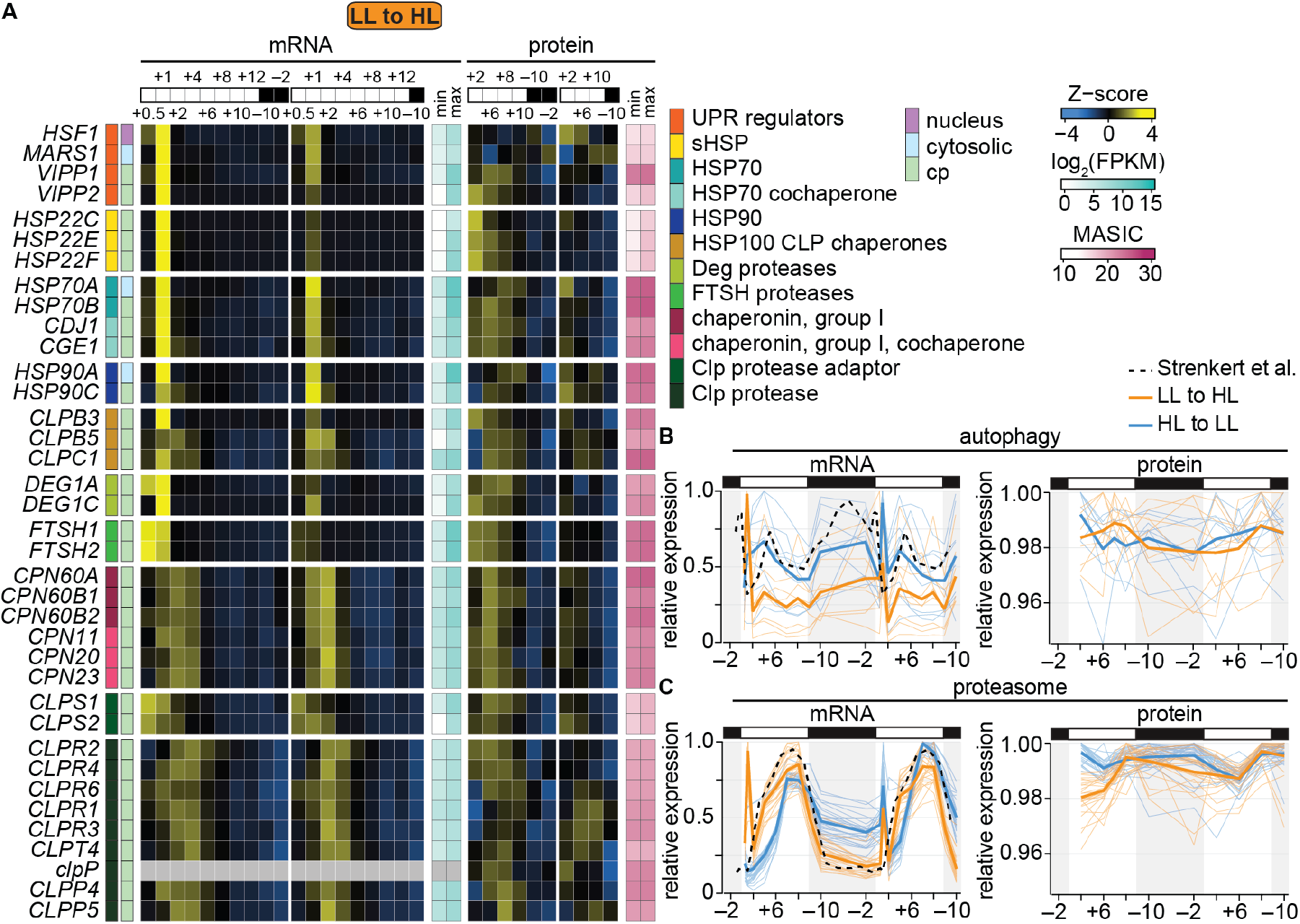
Chaperones, proteases, and the autophagy pathway are rapidly induced in LL-acclimated cells transitioned to HL. (A) Changes in mRNAs and proteins involved in chloroplast protein homeostasis. Z-scores of mean mRNA abundances (FPKM) and Z-scores of mean protein abundances (MASIC value) are used to show patterns over time. Minimum and maximum FPKM and MASIC values are also shown to demonstrate the dynamic range. Protein localization is listed according to the review by Schroda and deVitry 2023. (B) Expression of autophagy genes during the diurnal transitions compared to that of ML-acclimated cells reported by Strenkert *et al*., 2019, represented as in Figure 5C–5D. (C) Expression of proteasome genes shown as in Figures 5C–5D.

Proteomics showed that the VIPPs, chaperones, and proteases were most abundant between the +2 and +8 timepoints of the first day of HL (Figure 6A), concomitant with the striking changes in chloroplast integrity (Figure 2). The chloroplast Group I chaperonins, their cochaperones, the Clp protease adaptor CLPS1, and the Clp monomers CLPR2 and CLPR4 also peaked in abundance during this period, although transcripts for these proteins accumulated more gradually over the first day of the transition rather than as a sharp induction at lights on.

Under photooxidative and proteotoxic stress in the chloroplast, Chlamydomonas has also been reported to induce autophagy-related genes (64–66). We observed that several autophagy genes were induced at lights on of the LL-to-HL transition (Figure 6B). We also observed a sharp induction of proteasome mRNAs at the beginning of the LL-to-HL transition, which was roughly 2X higher than that observed during the HL-to-LL transition and was absent in ML-acclimated cells (Figure 6C) (31). Although some of these mRNAs exhibited peaks in abundance in the second morning of the HL-to-LL transition and in ML-acclimated cells (31), the peaks were of substantially lower magnitude. For example, transcripts encoding ATG8, a protein critical for autophagosome formation that is often used as a marker for autophagy, increased to ∼1700 FPKM at the first +1 timepoint of the LL-to-HL transition but only ∼700 FPKM in the HL-to-LL transition.

The proteins involved in autophagy were not co-expressed, but given that many of them are degraded along with the autophagic cargo (67), this was not unexpected. The abundance of the proteasome subunits on the other hand increased gradually over the first HL day. Proteasome function feeds back on *de novo* protein synthesis and is necessary for efficient recovery from photoinhibition in Chlamydomonas (68). It has also been linked to chloroplast proteostasis (69). Taken together, rapid induction of chaperones, proteases, and the autophagy pathway could be important for repair and remodeling during the transition from diurnal LL to HL.

## Discussion

We have undertaken a systems analysis of synchronized Chlamydomonas populations acclimated to dim days and bright days as they adjust to an unanticipated diurnal light intensity. Chlamydomonas populations acclimated to diurnal LL exhibit robust growth in a surprise HL day, achieving similar biomass yields over their first day in HL as do populations acclimated to diurnal HL through weeks of exposure (Figure 1). This is despite severe photoinhibition and loss of chloroplast integrity that occur during their first day in HL (Figure 2). LL-acclimated cells can recover PSII efficiency by the end of their first HL day, and by their second day, they perform almost indistinguishably from cells that are long-term accustomed to this daylight intensity. In contrast, cells acclimated to diurnal HL suffer growth limitation and do not divide after a surprise LL day (Figure S2), but these cells also adjust their physiology to improve growth yields just one day later. Taken together, these results demonstrate that Chlamydomonas is resilient to challenging light intensities under diurnal cycles and is competent to acclimate to variation in daylight intensities within 48 h.

The synchronous populations and temporally resolved changes make this an excellent system for studying photoinhibition and repair. Microscopy of live and fixed Chlamydomonas cells showed that LL-acclimated cells decreased thylakoid membrane stacking within the first 6 h of HL (Figures 2). Decreased stacking may allow the PSII repair machinery to access damaged PSII in these cells, providing a plausible mechanism for the recovery in F_v_/F_m_ that we observed in the latter half of the day (17, 41). On the other hand, segregation of the two photosystems is thought to prevent excitation energy spillover (70). Thus, it is also possible that rapid unstacking may result in spillover midday and contribute to the decrease in PSII efficiency.

Unlike at other timepoints, the chloroplast stroma appeared less densely packed with ribosomes than the cytosol did at the first +6 timepoint in HL, suggesting that the chloroplasts were swollen (Figure 2). Chloroplast swelling has been documented in *Arabidopsis* cells during photoinhibition and apparent damage to the chloroplast envelope (46), and the swollen chloroplasts are cleared through microautophagy of the entire organelle (chlorophagy) (46, 47, 71). We found that the light stroma phenotype was largely resolved across the Chlamydomonas LL-to-HL population within just 4 h. Chlamydomonas does not possess a canonical microautophagy pathway, nor is it likely that the cell’s only chloroplast would be completely turned over in just 4 h. However, piecemeal macroautophagy could play a role in the recovery of chloroplast integrity. Autophagy genes and others enriched for the “vesicle-mediated transport” GO term were induced at the onset of the LL-to-HL transition (Figures 3 and 6), and they are induced under chloroplast proteotoxic stress in Chlamydomonas (64–66).

The LL-acclimated cells exhibited the transcriptional signature of the chloroplast unfolded protein response at the start of the transition (Figure 6). Several VIPP proteins, chaperones, and proteases accumulated midday. Overexpression of VIPP1 suppresses chloroplast swelling in Arabidopsis (46). The VIPP proteins interact with the chloroplast chaperone HSP70B in Chlamydomonas and were proposed to sense damaged or misfolded proteins in the thylakoid membranes, recruit chaperones and proteases to such sites, and contribute to membrane remodeling (52, 60, 72, 73). Future work should 1) confirm whether the light stroma phenotype in fact reflects chloroplast swelling, and 2) determine the contributions of autophagy, proteostasis pathways, and VIPP proteins in the rapid restoration of chloroplast form and function during a surprise HL day.

Among the proteases induced at the beginning of the diurnal LL-to-HL transition were FTSH1, FTSH2, and DEG1C (Figure 6). These proteases cooperatively degrade the D1 polypeptide (61, 62, 74). However, the PSII core subunits PsbA (D1), PsbD (D2), PsbC (CP43), and PsbB (CP47) increased during the first HL morning, while the abundance of many other photosystem subunits and LHC proteins decreased (Figure 5), suggesting net D1 synthesis. Since we prepared RNA-Seq libraries using poly-A selection, we did not capture the chloroplast-encoded PSII mRNAs. However, previous reports have shown that *psbA* transcription does increase upon a transition from dark to light, as does the degradation rate of *psbA* mRNAs and several other chloroplast-encoded mRNAs (75). We did observe an apparent depletion of nucleus-encoded PSII mRNAs at lights on, which was more dramatic in cells transitioned from LL to HL than those transitioned from HL to LL or maintained in diurnal ML (Figure 5) (31). mRNAs encoding LHCII, PSI, and LHCI proteins also appeared to be depleted at the onset of the first HL day, and their accumulation over the light phase was delayed relative to populations maintained in diurnal ML (Figure 5). HL-dependent decreases in transcription, translation, and half-life of *LHCB* mRNAs have been demonstrated in Chlamydomonas (13, 44, 76, 77). We hypothesize that the apparent depletion of nucleus-encoded *PSB, LHCBM, PSA*, and *LHCA* mRNAs reflects other such light-responsive decreases in mRNA half-life.

Dark-to-light transitions are known to induce *LHCSR* and *PSBS* expression in Chlamydomonas, and the magnitude of mRNA accumulation at this time depends on the light intensity (3, 31). Here, we have found that the level of induction also depends on the light intensity experienced the previous day (Figure 4). LL-to-HL cells had much higher levels of these mRNAs at the first dark-to-light transition, whereas HL-to-LL cells increased their induction on the second dark-to-light transition. This dependence on the prior day’s light environment could stem, in part, from differences in the light absorption capacity and resulting photosynthetic electron transfer. Inhibition of photosynthetic electron transfer by DCMU has been shown to suppress the induction of *LHCSR3A* (3, 26). Thus, HL-acclimated cells with low photosynthetic capacity that are surprised with LL at lights on may have lower induction than do cells that have increased their inventory of antenna complexes by the second LL day, and *vice versa* (Figure 5).

In our controlled, reductionist experimental system, the light intensity remains constant during the light phase. But in nature, a day may start off gloomy and turn out bright, or the fog may roll in after a sunny morning. Future work should interrogate Chlamydomonas’ competence to re-acclimate to new light intensities at various times of the diurnal cycle and/or cell cycle. Given our results demonstrating the alga’s impressive flexibility, we suspect that Chlamydomonas has evolved to stay light on its feet and can adapt to just about any day.

## Materials and Methods

### Strains and culture conditions

*Chlamydomonas reinhardtii* strain CC-5390 was grown in High Salt (HS) medium with a modified trace element solution aerated with filter-sterilized air in flat-panel turbidostat FMT 150 Photobioreactors (Photon Systems Instruments, Drásov, Czechia) as previously described (30, 31). Cultures were subjected to diurnal cycles of 12 h light (80% blue light, 20% red light) at 50 (LL) or 1000 (HL) μmol photons m^−2^ s^−1^ and 28°C followed by 12 h dark and 18°C. The LL-acclimated population was first synchronized in diurnal ML (200 µmol photons m^−2^ s^−1^) for 1 week and then acclimated to diurnal LL (50 µmol photons m^−2^ s^−1^) for > 1 week. The HL-acclimated population was synchronized and acclimated in diurnal HL (1000 µmol photons m^−2^ s^−1^) for > 1 week. At the start of the experiment (–12, t = 0 h), culture density was set to OD_680_ = 0.4 and medium flow was stopped. Then, the LL-acclimated populations were subjected to diurnal HL conditions, and the HL-acclimated populations were subjected to diurnal LL conditions. After the first 26 h (–10), experimental cultures were diluted back to OD_680_ = 0.4 and medium flow was stopped again before the second day of the experiment. Once a photoacclimated population had been transitioned to a new light intensity and used for a two-day experiment, it was discarded and replaced with a fresh, naïve culture. Experimental replicates refer to independent experiments performed on cultures grown in different bioreactors months apart from each other. All measurements were taken for at least three experimental replicates.

### Productivity measurements

Cell growth was assessed by continuously monitoring culture OD_680_, periodically sampling for cell number and diameter with a Z2 Coulter Particle Count and Size Analyzer (Beckman Coulter, CA, USA), and periodically measuring NPOC using a TOC-L Shimadzu Total Organic Carbon Analyzer (Shimadzu, Kyoto, Japan). Photosynthetic activity was assessed by measuring F_v_/F_m_, NPQ, and steady-state Chl fluorescence emission spectra at 77K. Cellular pigment content was measured by HPLC. Detailed methods for NPOC, Chl fluorescence, and pigment content measurements are available in SI Information.

### Cellular morphology

Cultures were sampled for live-cell imaging by Airyscan microscopy of Chl fluorescence. Samples were kept on ice in the dark for <30 min prior to sample preparation and imaging. At least 6 representative cells were imaged per condition, and one representative image was chosen for display. For TEM, cells were fixed in 2% glutaraldehyde in HS medium at 4 °C with rotatory agitation in the dark for >10 h. Cells were stained, dehydrated, embedded, sectioned, and imaged as previously described (30). At least 15 representative cells were imaged from each sample, and one representative image was chosen for display. More details on sample preparation and imaging can be found in SI Information.

### RNA-Seq

Total RNA was extracted at 19 timepoints using the ZymoBIOMICS RNA Mini Kit (Zymo, CA, USA). Total RNA was subjected to poly(A) selection, RNA-Seq library construction, and sequencing on the Illumina HiSeq 3000 platform by the University of California Los Angeles Technology Center for Genomics and Bioinformatics (UCLA, CA, USA) using standard kits and protocols (Illumina Inc. CA, USA). Reads were mapped to the *Chlamydomonas reinhardtii* reference genome assembly and annotations v6.1 (78).

8 of the 114 samples experienced changes in sample volume during sample shipment and were flagged by the sequencing facility for possible evaporation or cross-contamination between wells of the 96-well plate (Figure S6A). PCA of all samples showed that these samples were dissimilar from the other experimental replicates of the condition (Figure S6B). These 8 libraries were therefore discarded, leaving at least two experimental replicates for each treatment group. Only nucleus-encoded mRNAs that were detected with a maximum FPKM > 1 were analyzed (15,305 genes) (Dataset S2). Additional information about RNA-Seq methodology can be found in SI Information.

### TMT proteomics

Cultures were sampled for tandem mass tag (TMT) proteomics at 10 timepoints. Cells were washed in 10 mM Na-phosphate buffer (pH 7.0), and the washed cell suspensions were flash-frozen in liquid N_2_ and stored at –80 °C until further processing. Cells were lysed by bead-beating in 8 M urea, the lysate was neutralized, and proteins were digested with 1:50 (w/w) trypsin and 1:20 (w/w) Lys-C proteases. Further sample processing, TMT peptide labelling, LC-MS/MS data collection, and proteomics data processing were performed as previously described (30).

While 11,489 proteins were detected across the dataset, only proteins that were detected in at least two of the three experimental replicates at all timepoints in either population were analyzed (9,567 proteins total) (Dataset S2). Additional details about proteomics methodology can be found in SI Information.

### Transcriptomic and proteomic data analysis

Gene-wise normalization was performed for each transition independently (LL to HL or HL to LL). PCA of the transcriptome and proteome was performed using the R package *PCAtools* (v2.16.0) on mRNAs with a minimum average FPKM > 1 (7613 genes), and proteins with a maximum average MASIC that was over the limit of quantitation (MASIC > 12.6) (8922 proteins in the LL-to-HL transition, 8984 proteins in HL-to-LL transition). *k*-means clustering analysis was conducted using the R package *stats* (v4.4.0). First, mean transcript abundances (as FPKMs) and mean protein abundances (as MASIC values) were Z-score normalized across time in a given transition (LL to HL or HL to LL). Only genes whose expression was detected at both the mRNA and protein levels at all timepoints of the transition were included (8821 genes in LL to HL, and 8868 genes in HL to LL) (Dataset S2), and the kmeans function was applied with centers = 16 and iter.max = 1000. Each cluster of genes was tested for GO term enrichment using the enricher function from the R package *clusterProfiler* (v4.12.0) with pvalueCutoff = 0.05 and pAdjustMethod = “BH”. Representative enriched GO terms are displayed. The full list of significantly enriched GO terms is available as Dataset S3.

Line plots of mean gene expression and the 95% confidence interval across the experimental replicates were generated in the R package *ggplot2* (v3.5.1) using the mean_cl_boot function. Composite heatmaps of gene expression patterns were generated in the R package *ComplexHeatmaps* (v2.20.0) (79).

## Author Contributions

S.D., S.S.M., and M.I. conceived of the study and designed experiments. M.S.L, K.K.N., and S.S.M provided resources and oversight. S.D. conducted the experiments. S.D., J.L.C, G.H., and M.I. performed physiological measurements. S.D., V.Z., S.D.G., C.D.N., and S.O.P. measured gene expression. S.D. conducted data analysis and wrote the manuscript with input from all authors.

## Acknowledgments

RNA-Seq was performed at the UCLA Technology Center for Genomics and Bioinformatics. Airyscan microscopy was performed at the Molecular Imaging Center, UC Berkeley Cancer Research Laboratory (supported by NIH 1S10OD025063). We gratefully acknowledge Reena Zalpuri and Danielle Jorgens at the UC Berkeley Electron Microscopy Lab for sample preparation, and Charles Perrino for technical assistance. This work was supported by The Gordon and Betty Moore Foundation Symbiosis in Aquatic Systems Initiative Investigator Award GBMF9203 (https://doi.org/10.37807/GBMF9203) to S.S.M. Microscopy and HPLC were supported by the US Department of Energy (DOE) Office of Science through the Photosynthetic Systems program in the Office of Basic Energy Sciences funding to K.K.N. and M.I.. Proteomics was performed on a project award from the Environmental Molecular Sciences Laboratory, a DOE Office of Science User Facility sponsored by the Biological and Environmental Research program under Contract No. DE-AC05-76RL01830. V.Z. was supported by the National Science Foundation Graduate Research Fellowship Program. K.K.N. is an investigator of the Howard Hughes Medical Institute.

## Competing Interest Statement

The authors declare no competing financial interests.

## Notes

### Competing Interest Statement

The authors have declared no competing interest.

