## Supplemental Information for "Light on its feet: Acclimation to high and low diurnal light is flexible in *Chlamydomonas reinhardtii*"

**This PDF file includes:**

Supporting text
Figures S1 to S6
Legends for Datasets S1 to S3
SI References

**Other supporting materials for this manuscript include the following:**

Datasets S1 to S3

### Supporting Information Text

#### Supplementary Methods

**Study design.** The physiology of transitioned cells was compared to data on diurnal HL- and LL-acclimated cells published previously (1). The gene expression of transitioned cells was compared over the time of the transition and to one another (LL to HL vs. HL to LL). In addition, patterns of mRNA accumulation over time were often compared to data collected with high temporal resolution on ML-acclimated cells published previously (2). The gene expression of the transitioned cells was not compared to that of the HL- and LL- acclimated populations, as the contribution of technical differences in data collection for the two experiments (e.g., different temporal resolution) was too great to draw meaningful conclusions from such comparisons, even after sample-wise and/ or gene-wise normalization.

**Nonpurgeable organic carbon measurements.** Total nonpurgeable organic carbon (NPOC) of cells was measured using a TOC-L Shimadzu Total Organic Carbon Analyzer (Shimadzu, Kyoto, Japan). 10 ml of culture was collected by centrifugation at 1680 × *g* and 4 °C for 2 min, and the cell pellets were washed once in 10 mM Na-phosphate buffer (pH 7.0). Cell pellets were stored at –20 °C until analysis (<1 month). To digest cells, pellets were vortexed in the remaining buffer, HCl added to achieve 5 M acid concentration, and then incubated at 65 °C overnight with constant agitation. Cell digests were diluted with MilliQ $H_2O$ for a final HCl concentration of 27 mM. Samples were sparged to remove inorganic carbon; some organic carbon may be purged from the sample by this method, so we report “nonpurgeable” organic carbon.

**Chl fluorescence measurements.** The maximum quantum efficiency of PSII ($F_v/F_m$) of cells dark-adapted for 15 min was measured using an IMAG-MAX/L MAXI Imaging PAM system (Heinz Walz GmbH, Effeltrich, Germany) as previously described (1).

To determine NPQ capacity, 12 ml of culture was collected into 50 ml beakers and agitated at 180 rpm to prevent hypoxia. The cells were dark-adapted for 30 min and then illuminated with far-red light to induce State 1 for 10 min for light-phase cells or 5 min for dark-phase cells. Cells were deposited onto glass fiber prefilters (Millipore Sigma, MA, USA) using a syringe, and the filters were placed on Whatman paper moistened with HS medium to prevent drying. Chl fluorescence was measured over light curves using an IMAG-MAX/L MAXI Imaging PAM system with gain set to 15, damping set to 5, and measuring light intensity and frequency both set to 1. $F_v/F_m$ was measured using a saturating pulse with intensity of 10 and pulse width of 0.72 s. Actinic light treatments of 50, 180, 550, 970, and 1490 μmol photons $m^{-2}$ $s^{-1}$ for 5 min were administered and light-adapted $F_m'$ was measured after each. NPQ was calculated as $(F_m – F_m')/F_m'$, and the maximum NPQ for each sample is reported.

To determine the steady-state Chl fluorescence emission spectra at 77 K, 750 μl of culture was collected into glass tubes and immediately frozen by submerging in liquid $N_2$. Fluorescence spectra were collected using a FluoroMax-4 spectrophotometer (Horiba Scientific, Kyoto, Japan) as previously described (1), with excitation of 420 nm and 2 nm slit size, and emission measured for 0.1 s at each wavelength from 650 to 780 nm and a 4 nm slit size. Fluorescence was normalized relative to the maximum fluorescence (resulting from PSII, ~684 nm). PSI fluorescence relative to the PSII fluorescence is reported as the ratio between the fluorescence at the 711 nm peak and the fluorescence at the 684 nm peak.

**Pigment content.** Cellular pigment content was measured by HPLC. 10 ml of culture was collected by centrifugation at 2500 × *g* for 2 min at 4 °C, and the pellets were flash-frozen in liquid $N_2$ and stored at –80 °C until processing. Pellets were thawed on ice, vortexed, and then the pellet was collected by centrifugation at 1000 × *g* for 3 min at 4 °C. 100 μl of 100% (v/v) ethanol was used to extract pigments from each pellet by vortexing for 3 min. The supernatant was collected by centrifugation at 21,000 × *g* for 5 min at ambient temperature. Another 50 μl of 100% (v/v) ethanol was added to the pellet to extract remaining pigment by vortexing for 3 min. The additional supernatant was collected by centrifugation at 21,000 × *g* for 5 min at ambient temperature. The combined extracted pigments were passed through a 0.22 μm nylon filter by centrifugation at 2650 × *g* for 1 min at ambient temperature. HPLC analysis was performed according to Gupta *et al.* (2015) with modifications as follows (3). 20 µl of the filtered extracted pigments was injected to HPLC and separated on a C30 reverse-phase column (YMC Carotenoid, 250 × 4.6 mm I.D., S-5 µm) coupled to a C30 guard column (YMC America) using mobile phases consisting of (A) 98% (v/v) methanol, (B) 95% (v/v) methanol, and (C) 100% (v/v) methyl *t*-butyl ether. Pigments were eluted with a linear gradient from 100% A to 60:40% A:C at a flow rate of 1 ml/min for 20 min, immediately followed by a linear gradient from 60:40% B:C to 100% C at a flow rate of 1 ml/min for 2 min, and re-equilibrate to initial condition by 30 min. The column temperature was maintained at 20 °C. Pigments were detected by absorbance at 445 nm with a reference at 550 nm by a diode array detector. Standards for carotenoids (antheraxanthin, α-carotene, β-carotene, lutein, neoxanthin, violaxanthin, and zeaxanthin) were purchased from DHI Laboratory Products (Hørsholm, Denmark). The quantification of each pigment was performed against calibration curves measured using these standards. Pigment content was normalized to cell number.

**Live-cell imaging.** 22 ml of culture was placed on ice in the dark for <30 min prior to sample preparation and imaging. Cells were collected by centrifugation at 1800 × *g* for 1 min and then resuspended in 0.7% (w/v) low-melting-point agarose in HS medium. Airyscan imaging of Chl fluorescence was performed using a Zeiss LSM 980 microscope equipped with an Airyscan detector and a Zeiss Plan-Apochromat 63×/1.4 NA DIC M27 Oil objective (Zeiss, Oberkochen, Germany). A 633 nm laser was used for excitation, and fluorescence emission was acquired through a 645 nm long-pass filter. At least 6 representative cells were imaged per condition, and one representative image was chosen for display. Image acquisition and analysis were performed using ZEN software (Zeiss, Oberkochen, Germany) and FIJI image analysis software (4, 5).

**Transmission electron microscopy (TEM).** 40 ml of culture was collected by centrifugation at 1800 × *g* for 1 min, and cells were fixed in 2% glutaraldehyde in HS medium at 4 °C with rotatory agitation in the dark for >10 h. Cells were stained, dehydrated, embedded, and sectioned as previously described (1). Micrographs were collected using a Technai 12 120 kV TEM with a UltraScan 1000XP CCD Camera (Gatan Inc, CA, USA). At least 15 representative cells were imaged from each sample, and one representative image was chosen for display.

**RNA extraction and sequencing.** Total RNA was extracted at 19 timepoints from 3 experimental replicates using the ZymoBIOMICS RNA Mini Kit (Zymo, CA, USA). 17 ml of culture was collected by centrifugation at 2500 × *g* for 2 min at 4°C, cell pellets were resuspended in 750 µl DNA/RNA Shield and transferred to ZR BashingBead Lysis Tubes containing 0.1 mm and 0.5 mm beads. Samples were immediately subjected to bead beating for 2 min in a Mini-BeadBeater 16 Cell Disrupter (BioSpec Products, OK, USA) in a 4 °C room. Then 400 µl supernatant was collected by centrifugation at 10,000 × *g* for 30 s and transferred to a nuclease-free tube. RNA was extracted according to the manufacturer's instructions, was flash frozen in liquid $N_2$, and stored at –80 °C.

Total RNA was subjected to poly(A) selection, RNA-Seq library construction, and sequencing on the Illumina HiSeq 3000 platform by the University of California Los Angeles Technology Center for Genomics and Bioinformatics (UCLA, CA, USA) using standard kits and protocols (Illumina Inc. CA, USA).

Reads from the 114 libraries were mapped to the *Chlamydomonas reinhardtii* reference genome assembly and annotations v6.1 (6) using STAR aligner (v2.7.9a) (7). The --alignIntronMax was set to 3000, allowing intron lengths up to 3000 base pairs. For multi-mapping reads, --outMultimapperOrder was set to Random, such that reads mapping to multiple locations were assigned randomly to the loci. The --outSAMmultNmax 1 parameter was applied to limit reporting to a single alignment per multi-mapping read.

mRNA abundance was quantified using the featureCounts function from the R package *Rsubread* (v2.1) (8), which assigns reads to genomic features. The countMultiMappingReads = TRUE option was used to include reads mapping to multiple locations. The allowMultiOverlap = TRUE option allows reads to be assigned to multiple overlapping features, and the fraction = TRUE option was applied to assign fractional counts to reads that overlap multiple features. The isPairedEnd = TRUE and countReadPairs = TRUE options were applied to ensure accurate counting of read pairs. DESeq2 (v1.45.0) was used for further data processing and normalization (9). Raw count data was imported and rounded to the nearest integer. Gene lengths were added to the DESeq2 dataset. A DESeqDataSet was created using DESeqDataSetFromMatrix() with a design formula of ~1. Gene lengths were incorporated using mcols(), and size factors were estimated with estimateSizeFactors(). FPKM values were calculated using the fpkm() function.

**TMT proteomics.** Cell pellets were collected for tandem mass tag (TMT) proteomics at 10 timepoints from three experimental replicates. 20 ml of culture was collected by centrifugation at 2500 × *g* at 4 °C for 2 min and washed in 40 ml 10 mM Na-phosphate buffer (pH 7.0). Cell pellets were resuspended in 500 µl 10 mM Na-phosphate buffer and transferred completely to 2 ml Eppendorf SafeLock tubes. Cells were collected by centrifugation at 16,000 × *g* at 4 °C for 20 s, then cell pellets were resuspended in 200 µl 10 mM Na-phosphate buffer by vortexing before being flash-frozen in liquid $N_2$. Samples were stored at –80 °C until further processing.

8 M urea was added to the cell pellets and cells were lysed by bead beating for 45 s at 5.5 m/s in a Bead Ruptor Elite bead mill homogenizer (OMNI International, GA, USA). Cell lysate was collected by centrifugation at 1000 × *g* at 4 °C for 10 min, and the protein concentration was determined by bicinchoninic acid (BCA) assay (Thermo Scientific, MA, USA). Samples were incubated with 10 mM dithiothreitol (DTT) at 60 °C for 30 min with constant agitation, followed by incubation with 40 mM iodoacetamide at ambient temperature for 30 min in the dark. Proteins were digested with 1:50 (w/w) Promega Platinum Trypsin (Promega, WI, USA) and 1:20 (w/w) Lys-C (FUJIFILM Biosciences, CA, USA) at 37 °C for 16 h, followed by cooling to 4 °C with constant agitation. Further sample processing, TMT peptide labelling, LC-MS/MS data collection, and proteomics data processing were performed as previously described (1).

### Supplementary Figures

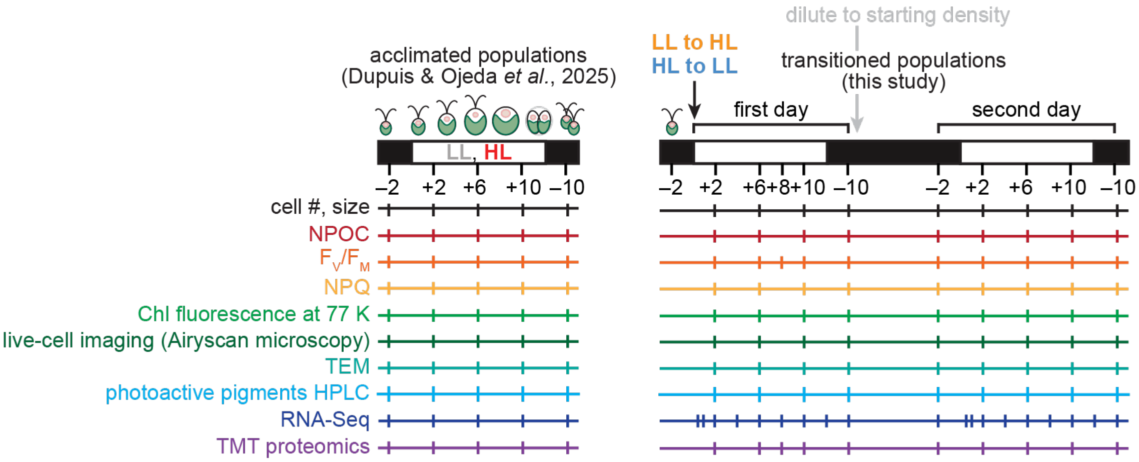


**Fig. S1. Measurements performed on acclimated populations (LL and HL) (1) and transitioned populations (LL to HL and HL to LL) over the diurnal cycle.** The black arrow indicates the time at which LL-acclimated populations were transitioned to diurnal HL and HL-acclimated populations were transitioned to diurnal LL (+0). The grey arrow indicates when the cultures were diluted to $OD_{680}$ of 0.4 prior to the second day of the experiment. Tick marks represent sampling timepoints.

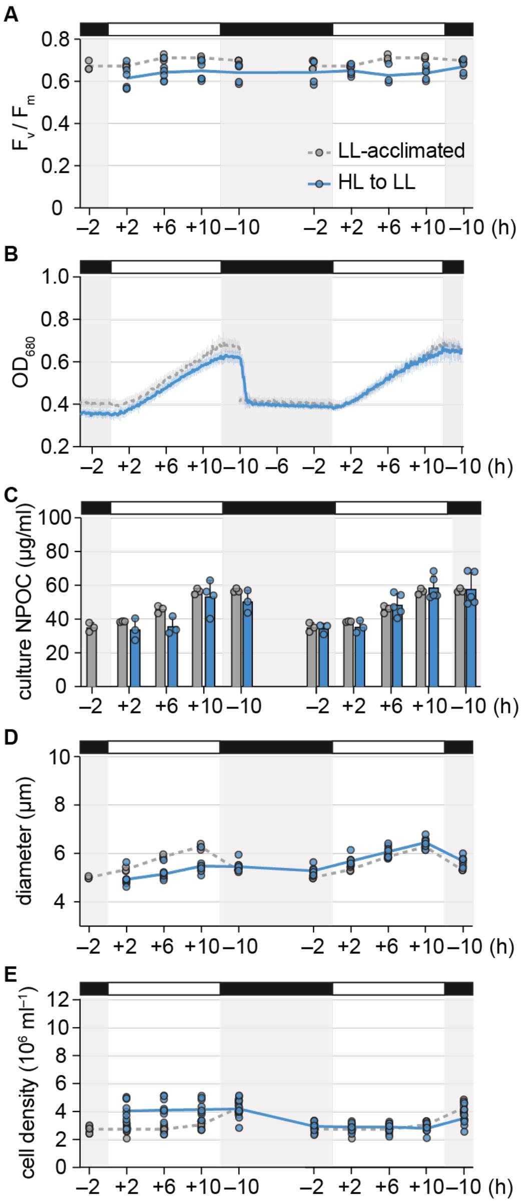


**Fig. S2. Acclimation to diurnal HL decreases cell growth and division upon a transition to LL.** The physiology of HL-acclimated cells transitioned to diurnal LL (blue) were monitored for two diurnal cycles. Results for LL-acclimated populations (grey) are reproduced from Dupuis & Ojeda *et al.*, 2025 (1).

(A) $F_v/F_m$.
(B) $OD_{680}$ monitored continuously. Error bars represent the standard deviation from the mean.
(C) NPOC concentration. Error bars represent the standard deviation from the mean.
(D) Mean cell diameter.
(E) Cell density.

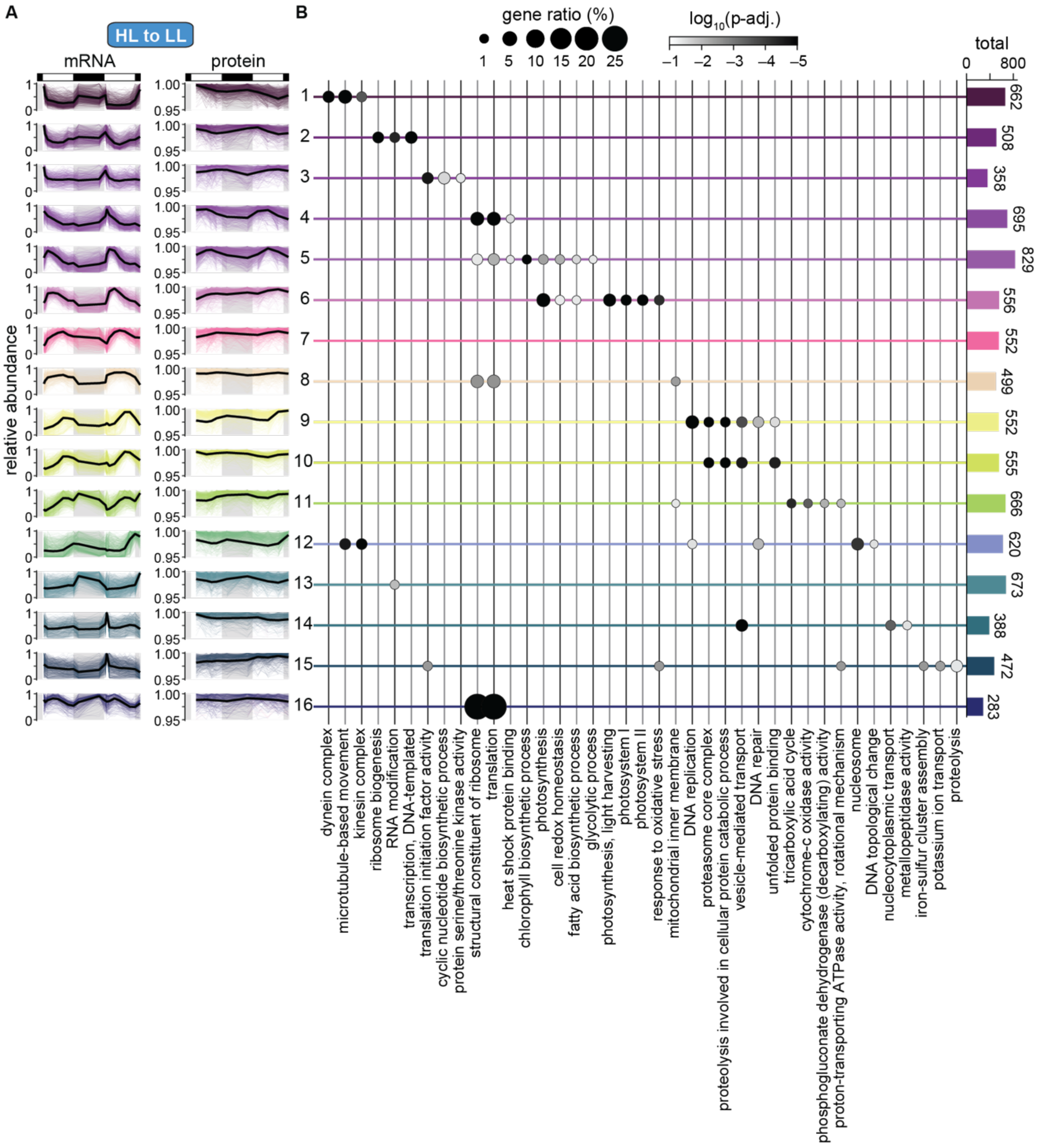


**Fig. S3. Although some mRNAs begin the HL-to-LL transition at already high abundance, new induction at lights on is not evident until the second day in LL.**

(A) Pattern of mRNA (left) and protein (right) accumulation for 16 distinct clusters of co-expressing genes during the HL-to-LL transition. Abundance was normalized relative to the maximum for each gene during the HL-to-LL transition. Colored lines show the relative abundance of individual mRNAs and proteins, while the black line shows the mean relative abundance. Cluster assignment is available in Dataset S2.

(B) Enrichment of representative GO terms in the 16 clusters of genes. Bars at the right show the total number of genes in each cluster. The full list of significantly enriched GO terms is available in Dataset S3.

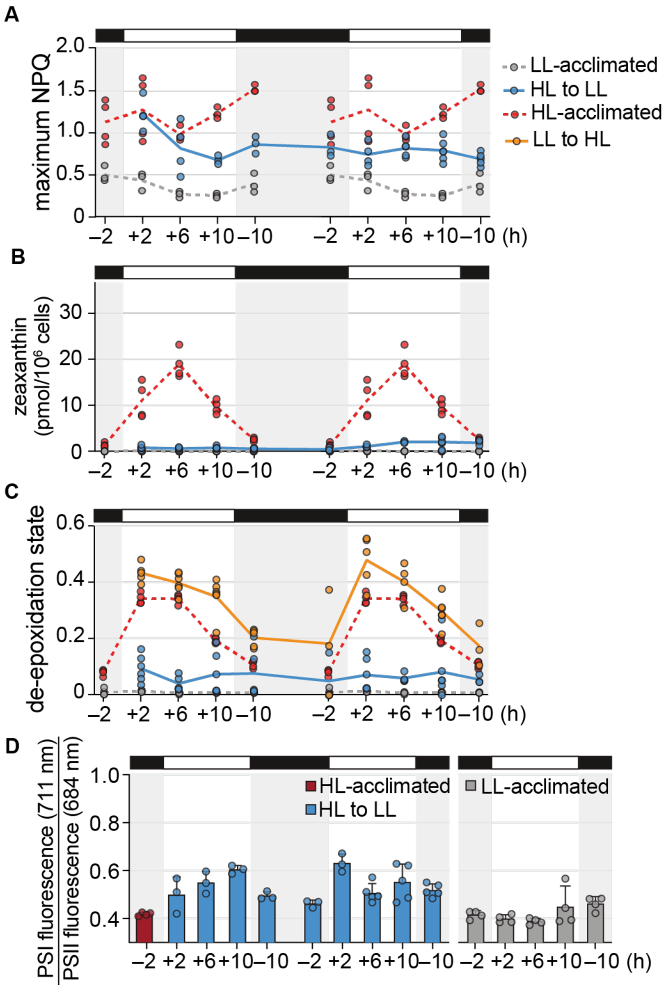


**Fig. S4. Photoprotective quenching is reduced during the transition from diurnal HL to LL.** Results for HL-acclimated populations (red) and LL-acclimated populations (grey) are reproduced from Dupuis & Ojeda et al., 2025 (1). Abundance of other photoprotection transcripts during the diurnal LL to HL transition.

(A) Maximum NPQ measured over 50–1500 µmol photons $m^{-2}$ $s^{-1}$ at each timepoint.
(B) Cellular zeaxanthin content.
(C) De-epoxidation state of xanthophylls, estimated as (0.5 antheraxanthin + zeaxanthin) / (violaxanthin + antheraxanthin + zeaxanthin).
(D) Relative Chl fluorescence from PSI (at 711 nm) to PSII (at 684 nm) measured at 77 K.

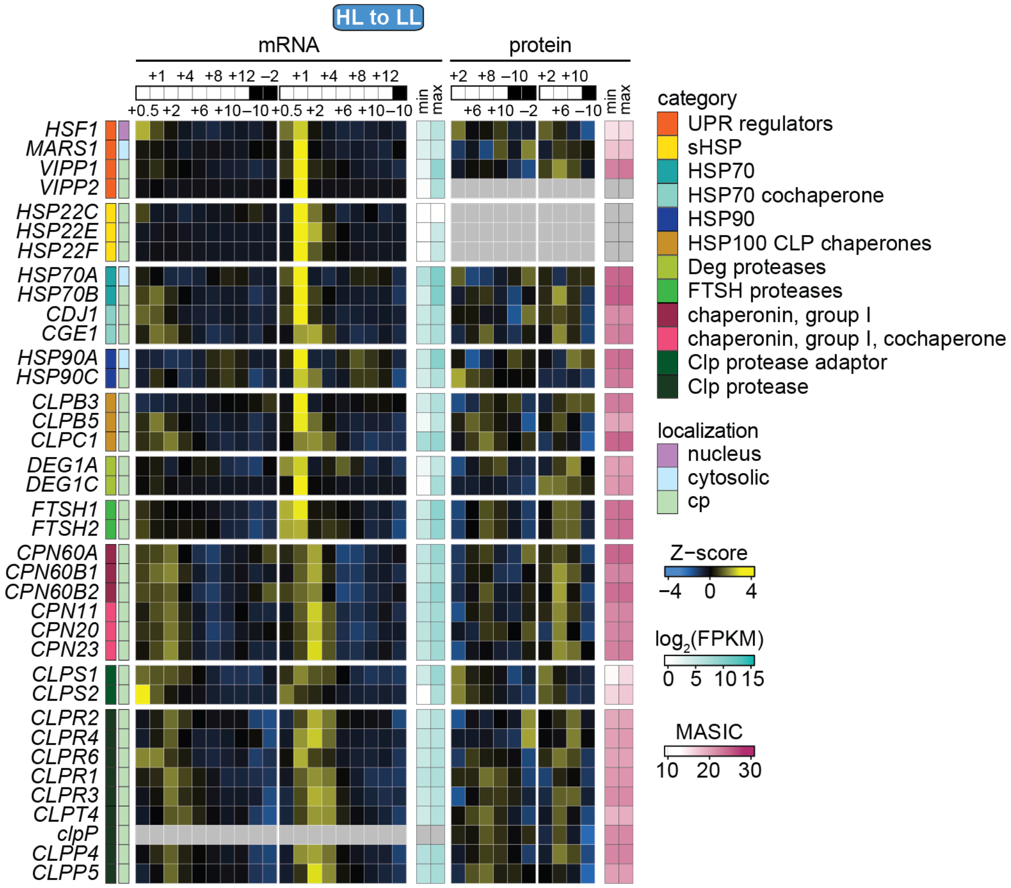


**Fig. S5. Induction of chloroplast chaperones, proteases, and their regulators is lower in the HL-to-LL transition than in the LL-to-HL transition.**

(A) Changes in mRNAs and proteins involved in chloroplast protein homeostasis. Z-scores of mean mRNA abundances (FPKM) and Z-scores of mean protein abundances (MASIC value) across 3 experimental replicates are used to show patterns over time. Minimum and maximum FPKM and MASIC values are also shown to demonstrate the dynamic range. Protein localization is listed according to the review by Schroda & de Vitry, 2023 (10).

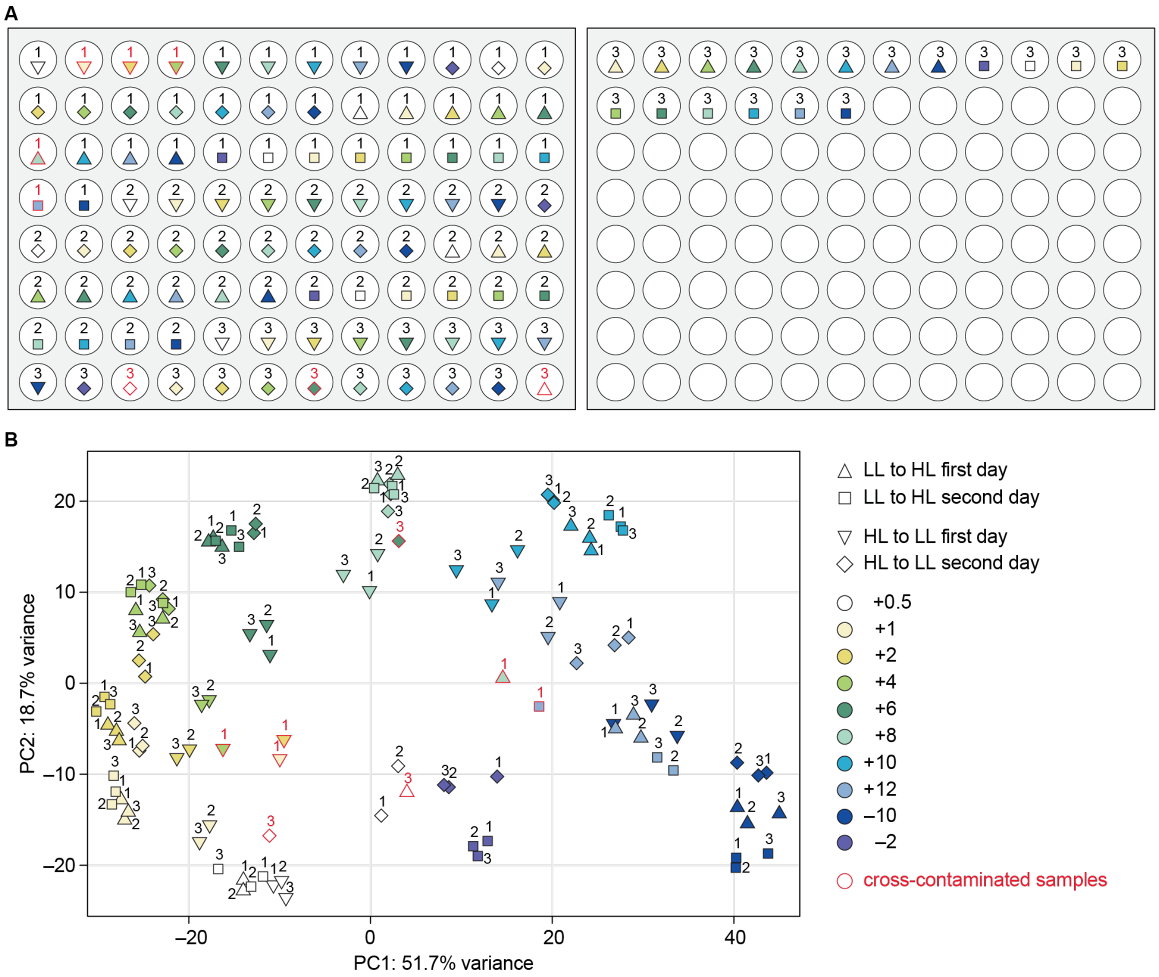


**Fig. S6. Several RNA samples were discarded due to sample cross-contamination.**

(A) Diagram showing how total RNA samples were arranged in 96-well plates for shipment to the sequencing facility. The numbers indicate the experimental replicate, the shapes indicate the day of the transition, and the symbol colors indicate the time of day. Samples outlined in red experienced changes in sample volume during shipment, indicating either sample evaporation or cross-contamination between wells.

(B) PCA of the abundance of the top 1000 most variable mRNAs from the experimental replicate samples indicated as in (A). The PCA positioned the samples that experienced changes in volume during shipment (red outlines) further away from the other replicates of the same timepoint in the transitions, providing evidence that these samples had been compromised. Data from these samples were therefore omitted. Nevertheless, all timepoints were still represented by at least two experimental replicates.

**Dataset S1 (separate file). Sample metadata for the transcriptome and proteome; related to Dataset S2.** This dataset contains metadata for the 106 samples that were analyzed by RNA-Seq and the 60 samples that were analyzed by TMT proteomics. It includes each sample's acclimation light intensity, experiment light intensity, timepoint, and replicate number. Additional details are provided in Materials and Methods and the SI Appendix.

**Dataset S2 (separate file). Genome-wide mRNA abundance (FPKM) and protein abundance (MASIC value) for Chlamydomonas populations transitioned from diurnal LL to HL or diurnal HL to LL; related to Figures 3, 4, 5, 6, S1, S3, S4, S5, and S6.** This dataset contains FPKM values for 15,305 nucleus-encoded mRNAs that were detected with a maximum FPKM above 1, and MASIC values for 9,567 proteins that were detected in at least two of the three experimental replicates of either the LL-to-HL population or the HL-to-LL population. "NA" indicates that the molecule was not detected in the sample. Gene symbols and functional descriptions from the *C. reinhardtii* CC-4532 v6.1 genome are included. *k*means cluster assignment for included genes is also provided. Metadata for each sample are available as Dataset S1. Additional details about the analyses are provided in Materials and Methods and the SI Appendix.

**Dataset S3 (separate file). Gene ontology enrichment of the *k*means clusters; related to Figures 3 and S3.** This dataset contains the complete results of the gene ontology (GO) enrichment analysis on the 16 clusters of genes shown in Figure 3E (LL to HL) and the 16 clusters of genes shown in Figure S3A (HL to LL). The analysis is described in the Materials and Methods.
